# The draft genome sequence of mandrill (*Mandrillus sphinx*)

**DOI:** 10.1101/367870

**Authors:** Ye Yin, Ting Yang, Huan Liu, Ziheng Huang, Yaolei Zhang, Yue Song, Wenliang Wang, Karsten Kristiansen

**Author notes:** These authors contributed equally to this work (Ye Yin, Ting Yang).

## Abstract

**Background:** Mandrill (*Mandrillus sphinx*) is a primate species which belong to Old World monkey (Cercopithecidae) family. It is closely related to human, serving as model for some human diseases researches. However, genetic researches and genomic resources of mandrill were limited, especially comparing to other primate species.

**Findings:** Here we sequenced 284 Gb data, providing 96-fold coverage (considering the estimate genome size of 2.9 Gb), to construct a reference genome for mandrill. The assembled draft genome was 2.79 Gb with contig N50 of 20.48 Kb and scaffold N50 of 3.56 Mb. We annotated the mandrill genome to find 43.83% repeat elements, as well as 21,906 protein coding genes. We found good quality of the draft genome and gene annotation by BUSCO analysis which revealed 98% coverage of the BUSCOs.

**Conclusions:** We established the first draft genome sequence of mandrill, which is valuable resource for future evolutionary and human diseases studies.

## Background Information

Mandrill (here, specifically referred to Mandrillus sphinx) is a primate species living in Africa. It is a large monkey specie belonging to Papionini tribe. It has been used as experimental models for human disease studies considering that the species of chimpanzee and gorilla despite that they are more closely related human [1], they are endangered species. Especially, mandrill has been used in immune systems researches such as bacterial infections [2, 3], parasite infections [4, 5] and viral diseases especially for the study of SIV strains infection [6, 7]. However, comparing to chimpanzee and gorilla, mandrill only has the mitochondrial genomes established, without comprehensive genome information available.

In this study, we established a draft genome sequence of mandrill using second generation sequencing technology, making it possible for future comparative genomic studies and human disease studies.

## Methods

### Sample preparation

We obtained 5 mL blood from the left jugular vein of an eighteen-year-old male mandrill from Beijing Zoo. The blood was collected to a plastic collection tube with 4% (w/v) sodium citrate, then snap frozen in liquid nitrogen and stored at −80 ˚C. Genomic DNA was extracted using the AXYGEN Blood and Tissue Extraction Kit according to the manufacturer’s instructions. To assess the quality, the extracted DNA was subjected to electrophoresis in 2% agarose gel and stained with ethidium bromide. DNA concentration was detected by Quant-iT™ PicoGreen ^®^ dsDNA Reagent and Kits (Thermo Fisher Scientific, USA) according to the manufacturer’s protocol.

### Library construction and sequencing

DNA of mandrill was used for library construction, according to protocols following descriptions in the previous publication [8]. A total of 7 libraries were constructed, and then sequencing was carried out on Illumina sequencer HiSeq2000. Within the seven libraries, three were short insert size libraries including insert sizes of 250 bp (sequenced to 150 bp in two ends), 500 bp and 800 bp (sequenced to 100 bp in two ends), respectively. The other four libraries were mate-pair libraries with insert sizes of 2 kb, 5 kb, 10 kb and 20 kb (sequenced to 90 bp in two ends). Then, using SOAPnuck[9], to filtered reads according to the criterions: i) reads with more than 10% Ns (ambiguous bases); ii) reads with more than 40% of low quality bases (quality score less than 10); iii) reads contaminated by adaptor (adaptor matched 50% with no more than one base mismatch) and PCR duplicated reads (identical reads in both ends).

### Genome assembly

In order to assess the genome features, 17-mers (17 bp sub-sequences) were extracted and subjected to the K-mer analysis. The reads from 250 bp, 500 bp and 800 bp insert libraries were used for this analysis.

Then the genome was assembled by short-reads assembly software SOAPdenovo2 [10] using the filtered data (with parameter settings pregraph-K 35; contig -M 1; scaff). Gaps were filled using paired sequence data from 3 libraries (250, 500, and 800 bp) with -p 31 parameters by GapCloser.

### Transposable elements and repetitive DNA

We utilized RepeatMasker v4.0.5 [11] and Repeat-ProteinMask to scan the whole genome for known transposable elements in the RepBase library v20.04 [12]. Then, RepeatMasker was then used again for identifying *de novo* repeat based on the custom TE library constructed by combining results of RepeatModeler v1.0.8 (RepeatModeler,RRID:SCR_015027) and LTR_FINDER v1.0.6 [13]. Prediction of tandem repeats was also done using Tandem Repeat Finder v4.0.7 [14] with the following setting: Match = 2, Mismatch = 7, Delta = 7, PM = 80, PI = 10, Minscore = 50, MaxPeriod = 2000.

### Protein-coding gene and non-coding RNAs annotation

After masked known TE repeat elements, genes were predicted using three categories of methods, including homolog based, evidence based and ab initio prediction. For homolog based annotation, protein sequences of *Macaca mulatta* (Ensemble 73 release), *Pan troglodytes* (Ensemble 73 release), *Nomascus leucogenys, Pongo abelii* (Ensemble 73 release), *Gorilla gorilla* (Ensemble 73 release), *Homo sapiens* (Ensemble 73 release) and were aligned to the mandrill genome using BLAT [15]. Then GeneWise [16] (Version 2.2.0) was used for further precise alignment and gene structure prediction. For evidence based prediction, 85,496 EST sequences were downloaded from NCBI with key words: *Papio anubis* cDNA and were aligned to the genome using PASA [17] for spliced alignments and assembly to detected gene structure. For ab initio prediction, we employed AUGUSTUS [18] (Version 3.1) to predict gene models in the repeat masked genome. Finally, these gene prediction results were combined using GLEAN [19]. In order to identify the function of the final gene set, three databases (SwissProt, KEGG, and TrEMBL databases) were searched for best matches using BLASTP (version 2.2.26) with an E-value of 1e-5. InterProScan software [20], which searches Pfam, PRINTS, ProDom and SMART databases for known motifs and domains, was also used for the gene function annotation.

To identify transfer ribonucleic acids (tRNAs), tRNAscan [21] was used. While for ribosomal ribonucleic acids (rRNAs) identification, 757,441 rRNAs from public domain were used to search against the genome with command -p blastn -e 1e-5. To identify RNA genes and other non-coding RNA (ncRNA), Rfam database [22] was used to search against the genome.

### Gene family clustering

Protein sequences of 11 species including *Gorilla gorilla, Homo sapiens, Macaca mulatta, Microcebus murinus, Nomascus leucogenys, Otolemur garnettii, Pan troglodytes, Pongo abelii, Tarsius syricht, Callithrix jacchus*, and *Mus musculus* were used to cluster the gene families. TreeFam (http://www.treefam.org/) was used to defined gene families in *Mandrillus sphinx*. Firstly, all-versus-all blastp with the e-value cutoff of 1e-7 for 12 species and then the possible blast matches were joined together by an in-house program. Thirdly, we removed genes with aligned proportion less than 33% and converted bit score to percent score. Finally, hcluster_sg (Version0.5.0, https://pypi.python.org/pypi/hcluster) was used to cluster genes into gene families.

### Phylogenetic tree construction

With gene family clusters defined, the fourfold degenerate (4D) sites of 5,133 single-copy orthologous among the 12 species were extracted for the phylogenetic tree construction. PhyML package [23] was used to build the phylogenetic tree with maximum-likelihood methods and GTR+gamma as amino acid model (1,000 rapid bootstrap replicates conducted). Based on the phylogenetic tree, divergence times of these species were estimated by using MCMCTree (http://abacus.gene.ucl.ac.uk/software/paml.html) with the default parameters. To further calibrate the evolution time in the tree, six fossil dates collected from the TimeTree database (http://www.timetree.org/) were used, including the divergence time between *Mus musculus* and human to be 85–93 million years ago (MYA) [24], divergent time between human and chimpanzee, gorilla, to be 6 MYA (with a range of 5–7) [25] and 9 MYA (range, 8–10) [26].

With the gene family clustering result, contraction and expansion can be detected to figure out the dynamic evolutionary changes along the phylogenetic tree. According to the phylogenetic tree and divergence time, CAFÉ [27] was used for gene family contraction and expansion analysis.

Demography was estimated using the pairwise sequentially Markovian coalescent (PSMC) model with the following setting: -N25 –t15 –r5 -p “4+25*2+4+6”. In order to scale the results to real time, 10 years per generation and a neutral mutation rate of 2.5e-08 per generation were used.

### Gene family expansions/contractions and positively selected genes

The selection pressure in mandrill were measured by comparing nonsynonymous (dN) and synonymous (dS) substitution rates on protein-encoding genes. This ratio would be equal to 1 if the whole coding sequence evolves neutrally. When dN/dS < 1, it’s under constraint, and when dN/dS > 1 it should be under positive selection. Positively selected genes (PSGs) was detected using models in the program package PAML version 3.14, neutral (M1 and M7) and selection (M2 and M8) models were used.

## Data Description

### Genome sequencing and assembly

Whole genome sequencing of mandrill yielded 426 Gb raw sequence data (146× considering the genome size of 2.9 Gb). After filtering, clean reads amounting to 289 Gb were obtained for genome assembly, comprising ∼71x from paired-end libraries and ∼26x from mate-pair libraries (**Table S1**). 212 Gb data were used for k-mer analysis, which resulted in the distribution of depth-frequency (**Figure S1**), with a secondary peak at half of the major peak coverage of ∼31×. In this way, the genome size of mandrill was estimated to be 2.90 Gb with notable heterozygosity. All clean data were used to generate the draft genome assembly, followed by gap filling. The assembled genome was 2.88 Gb (covered 99.31% of estimated genome size) in length. The contig N50 was 20.5 kb with longest contig to be 211 kb, and the scaffold N50 was 3.6 Mb with the longest scaffold to be 19.1 Mb. 634 longest scaffolds consisted more than 80% of the whole genome (**Table 1**).

**Table 1.** Summary of the mandrill genome assembly.

|  | **Contig** |  | **Scaffold** |  |
| --- | --- | --- | --- | --- |
|  | Size (bp) | Number | Size (bp) | Number |
| **N90** | 5,266 | 141,475 | 638,217 | 936 |
| **N80** | 9,025 | 101,618 | 1,303,160 | 634 |
| **N70** | 12,638 | 75,505 | 1,962,294 | 457 |
| **N60** | 16,336 | 56,061 | 2,730,696 | 332 |
| **N50** | 20,483 | 40,751 | 3,564,730 | 241 |
| **Longest** | 211,017 | ---- | 19,105,867 | ---- |
| **Total size** | 2,798,997,503 | ---- | 2,882,689,325 | ---- |
| **Total number ($\geq 100$ bp)** | ---- | 455,069 | ---- | 215,140 |
| **Total number ($\geq 2$ kb)** | ---- | 194,923 | ---- | 4,742 |

### Genome annotation

Using a combination of *de novo* and homology-based methods, about 42.22% (mandrill) of the assembled genome were transposable elements (TEs) (**Table S2**). Long Interspersed Nuclear Elements (LINEs) were less in mandrill genome (∼17%) comparing to human genome (∼21%), while Short Interspersed Nuclear Elements (SINEs) were similar (∼12%), especially the *Alu* elements to have quite similar proportion (10%∼11%) **(Table S3)**, reflecting that the *Alu* elements were the conserved within primate genomes as previously described [28].

The gene set of the mandrill genome contains 21,906 protein-coding genes (**Table S4**), the average gene length was 39,087 bp with the average intron length of 5,785 bp. 21,622 (98.70%) of the predicted genes functionally annotated (**Table S5**). Three types of ncRNAs were annotated in mandrill genome, including tRNAs, rRNAs, and snRNAs. In total, 4,278 short noncoding RNA sequences were identified in the mandrill genome (**Table S6**).

The quality of mandrill genome and gene completeness were assessed by conducting the Benchmarking Universal Single-Copy Orthologs (BUSCO) analysis [29]. 98% of BUSCOs were completely detected in the assembled genome (2981: complete and single-copy; 170: complete and duplicated) among 3,023 tested BUSCOs. The numbers of fragmented and missing BUSCOs were 28 and 14, respectively (**Table S7**).

### Gene family and phylogenetic analysis

For mandrill, we identified 15,368 gene families with 1,387 genes not clustered among 12 species, and 87 families were found to be unique (**Table S8**). These unique gene families were significantly enriched in function annotation with GO:0006412 of translation (GO level: BP, P=6.29e-33), GO:0003735 of structural constituent of ribosome (GO level, BP, P=6.29e-33) (**Table S9**). Comparing to human (*Homo sapiens*), macaque (*Macaca mulatta*), gorilla (*Gorilla gorilla*) and marmoset (*Callithrix jacchus*), 627 gene families, with 1,293 genes, were found to be unique in the mandrill and 5,133 single-copy orthologous genes were found to be shared among all the 12 species (**Figure 1a)**.

**Figure 1.**
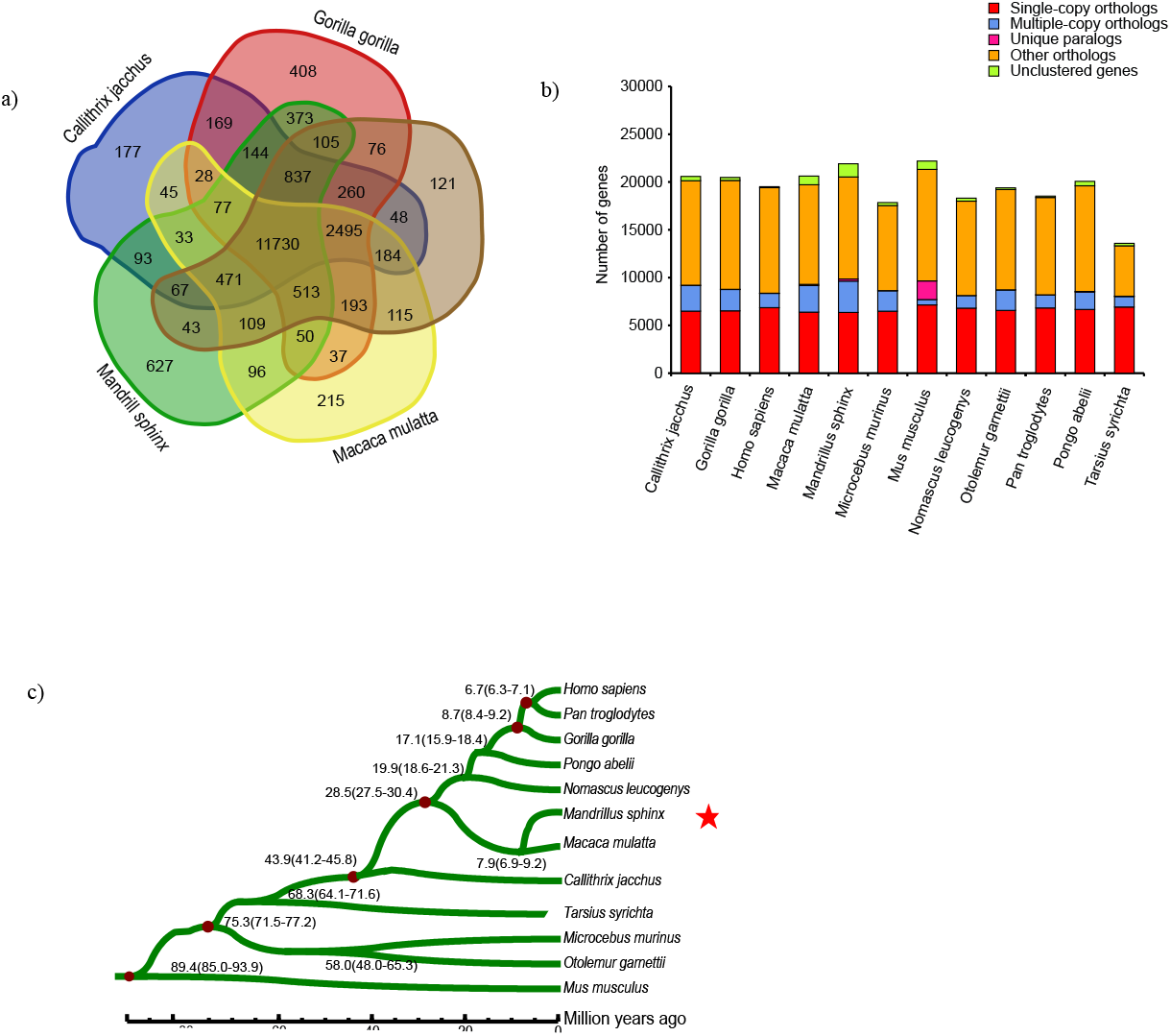
Evolution of mandrill in gene families and phylogeny. **a)** The Venn diagram of the gene families in the human (*Homo sapiens*), macaque (*Macaca mulatta*), gorilla (*Gorilla gorilla*), marmoset (*Callithrix jacchus*) and mandrill. **b)** Comparison of orthologous genes among 11 primates and mouse. **c)** The maximum-likelihood phylogenetic treebased onthe 4-fold degenerate sites of 5,133 single copy gene familiesin the 12 species.

The maximum-likelihood phylogenetic tree (**Figure 1b**) of the 12 species using 5,133 single-copy genes indicated that mandrill was located in the same clade with macaque and this clade diverged from human clade about 28.5 (27.5–30.4) MYA while the divergence time between Cercopithecoidea and Hominoidea was estimated to be 26.66 (24.29–28.95) MYA using mitochondrial genome sequences [30]. Mandrill were estimated to split from macaque about 7.9 (6.9–9.2) MYA which was different from the previous estimation which was 6.6 (6.0–8.0) MYA [31].

In mandrill lineage, there were 797 expanded and 3,982 contracted gene families (**Figure S2**). Expanded gene families were found to be significantly enriched in the functions of biosynthetic process, structural constituent of ribosome, nucleosomal DNA binding, G-protein coupled receptor activity, olfactory receptor activity, glucose catabolic process, peptidyl-prolyl isomerization, as well as carbon fixation in photosynthetic organisms and electron transport chain pathway (**Table S10**). In mandrill, peptidylprolyl isomerase A (PPIA) was significantly expanded (GO:0003755, P= 3.60E-89). The PPIA belongs to the peptidyl-prolyl cis-trans isomerase (PPIase) family which catalyze the cis-trans isomerization, folding of newly synthesized protein, combination of several transcription factors and regulating many biological processes including inflammation and apoptosis, even acting in cerebral hypoxia-ischemia. In stress environment when presence of reactive oxygen species (ROS), cell will secrete PPIA to induce an inflammatory response and mitigate tissue injury. The peroxiredoxin-6 (PRDX6) family, which can reduce peroxides and protection against oxidative injury during metabolism, was also significantly expanded (GO:0051920, P = 0.000641).

In total, 657 PSGs were identified with significant enrichment in the molecular functions of kinase activity, transferase activity, phosphotransferase activity and etc. (**Table S11**). Further investigating functions of these PSGs, 34 genes were found to be innate immunity response genes by searching InnateDB. Interactions of these genes were predicted by STRING: functional protein association networks (http://string-db.org/cgi). As shown in **Figure S3**, STAT1, IL5, IL1R1, ATG5, CREB1, DICER1, PIK3R1 genes may have important roles in immune system, which are strongly associated with stress resistance and wound healing. Finally, PSGs related to innate immunity response were found to be enriched in terms of GO:0080134: regulation of response to stress (GO level: BP, P=1.59e-05), GO:0006955: immune response (GO level: BP, P value=4.11e-05) and KEGG:4640: Hematopoietic cell lineage(P=6.83e-05) (**Table S12**).

The demographic history of a species reflects historical population changes thus would be important to understand from the genome. We inferred a noticeable population bottleneck in the demographic history of the mandrill (**Figure 2**). Around 28 thousand years (kys) ago, the mandrill population went through a sharp increase, followed by a noticeable bottleneck from a peak of 61,000 and 47,000 to ∼6,500 around 17 kys ago. The increase of population size was coincident with the increase of human population, probably indicating climate change suitable for increase of mammals, while the recent bottleneck of mandrill populations is different from the recent increase of the human population.

**Figure 2.**
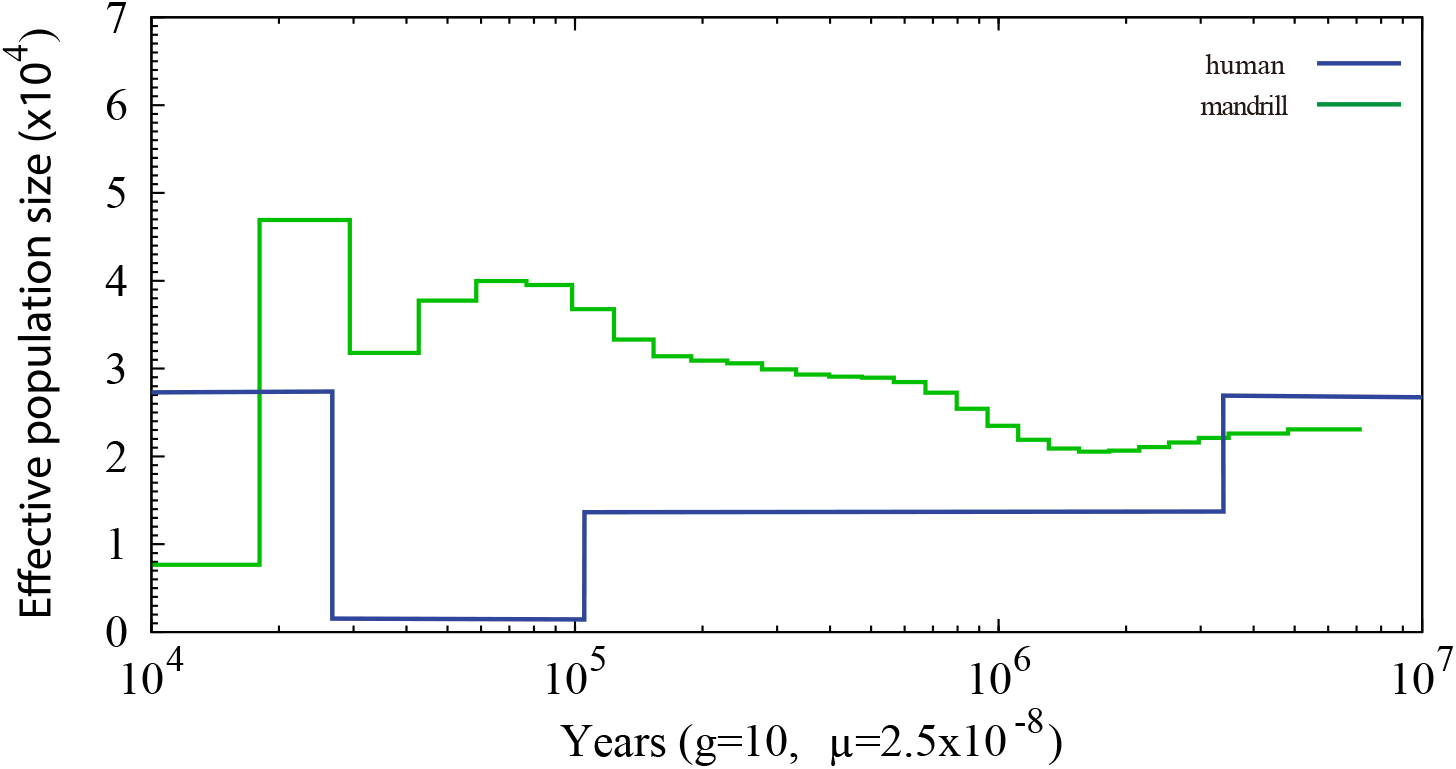
The demographic changes of mandrill comparing to human. The x-axis indicates the time (from left to right indicated from recent to ancient), and the y-axis indicates the estimated effective population size.

### Comparing the major histocompatibility complex (MHC) region

Major histocompatibility complex (MHC) is important for immune system which code a series of genes to assist cells recognizing foreign substances. Checking the assembled genomes of mandrill, relative complete assembly of MHC region was found on Chromosome 4. In order to check the assembly quality, reads were mapped back to the MHC class I region to show good coverage and pair-end/mate-pair relationships (**Figure S4**), supporting assembly of MHC region. Since the MHC region is highly repetitive, a detailed repeat annotation was carried out for both mandrill and human MHC class I regions (from gene *GABBR1* to gene *MICB* in the direction from the telomere side to the centromere side) with the same parameter to find similar repeat content for the two species in this region (48.27% in mandrill comparing to 51.03% in human) (**Table S13**). *HLA* genes are important for immune recognition thus HLA genes were further checked and compared to human. In MHC class I region of human, there were 50 genes in total including 6 HLA genes, while in mandrill MHC class I region, only 4 HLA genes were identified. Searching the whole genome other than the MHC region, another 4 HLA genes were identified, making the total number of HLA genes to be 8 in mandrill. However, further inspection of the 8 HLA genes in mandrill resulted in finding 5 of them harbored start or stop codon changes, prematurely terminated changes or frameshift mutations (**Figure 3**), reflecting the genetic mechanisms of differences in immune response between mandrill and human.

**Figure 3.**
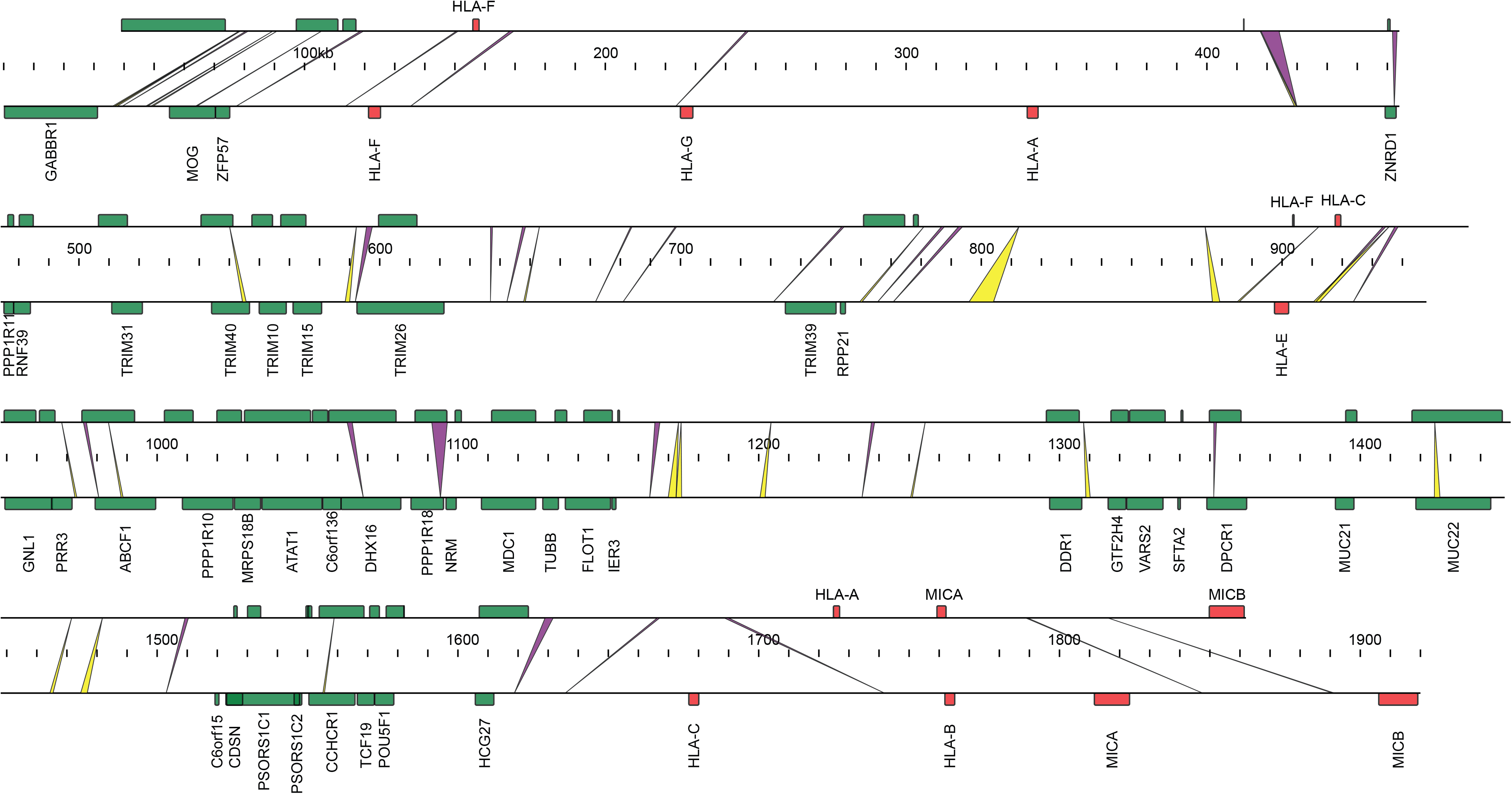
Synteny of MHC class I region between mandrill and human. Red boxes indicate the MHC genes. Green boxes indicate other protein coding genes. Yellow and green indicates deletions and insertions in mandrill respectively.

## Availability of supporting data

The raw sequencing data described in this article are available in the NCBI SRA repository, under the project number of PRJNA436891.

## Abbreviations

bp: base pair
kb: kilo base
Gb: giga base
MYA: million years ago
PSMC: pairwise sequentially Markovian coalescent
TE: transposable element
BUSCO: Benchmarking Universal Single-Copy Orthologs

## Ethics statement

Animal collection were approved by Beijing Zoo and utility were in accordance with guidelines from the China Council on Animal Care.

## Additional files

Additional file 1

Supplementary Figures – Figure S1 to Figure S4

Additional file 2

Supplementary Tables – From Table S1 to Table S13 (except Table S10 and Table S12)

Additional file 3

Table S10. GO term enrichment of gene families expanded in mandrill.

Additional file 4

Table S12. GO and KEGG enrichment of the positively selected genes related to innate immunity.

